# Trans-ethnic polygenic analysis supports genetic overlaps of lumbar disc degeneration with height, body mass index, and bone mineral density

**DOI:** 10.1101/316737

**Authors:** Xueya Zhou, Ching-Lung Cheung, Tatsuki Karasugi, Jaro Karppinen, Dino Samartzis, Yi-Hsiang Hsu, Timothy Shin-Heng Mak, You-Qiang Song, Kazuhiro Chiba, Yoshiharu Kawaguchi, Yan Li, Danny Chan, Kenneth Man-Chee Cheung, Shiro Ikegawa, Kathryn Song-Eng Cheah, Pak Chung Sham

## Abstract

Lumbar disc degeneration (LDD) is age-related break-down in the fibrocartilaginous joints between lumbar vertebrae. It is a major cause of low back pain and is conventionally assessed by magnetic resonance imaging (MRI). Like most other complex traits, LDD is likely polygenic and influenced by both genetic and environmental factors. However, genome-wide association studies (GWASs) of LDD have uncovered few susceptibility loci due to the limited sample size. Previous epidemiology studies of LDD also reported multiple heritable risk factors, including height, body mass index (BMI), bone mineral density (BMD), lipid levels, etc. Genetics can help elucidate causality between traits and suggest loci with pleiotropic effects. One such approach is polygenic score (PGS) which summarizes the effect of multiple variants by the summation of alleles weighted by estimated effects from GWAS. To investigate genetic overlaps of LDD and related heritable risk factors, we calculated the PGS of height, BMI, BMD and lipid levels in a Chinese population-based cohort with spine MRI examination and a Japanese case-control cohort of lumbar disc herniation (LDH) requiring surgery. Because most large-scale GWASs were done in European populations, PGS of corresponding traits were created using weights from European GWASs. We calibrated their prediction performance in independent Chinese samples, then tested associations with MRI-derived LDD scores and LDH affection status. The PGS of height, BMI, BMD and lipid levels were strongly associated with respective phenotypes in Chinese, although phenotype variances explained were lower than in Europeans which would reduce the power to detect genetic overlaps. Despite of this, the PGS of BMI and lumbar spine BMD were significantly associated with LDD scores; and the PGS of height was associated with the increased the liability of LDH. Furthermore, linkage disequilibrium score regression suggested that, osteoarthritis, another degenerative disorder that shares common features with LDD, also showed genetic correlations with height, BMI and BMD. The findings suggest a common key contribution of biomechanical stress to the pathogenesis of LDD and will direct the future search for pleiotropic genes.

## Introduction

Human intervertebral discs (IVDs) are fibrocartilaginous structures that lie between adjacent vertebrae. These IVDs hold the vertebrae together, facilitate some vertebral motion, and act as shock absorbers to accommodate biomechanical loads (Oxland 2016). IVD is composed of a gel-like nucleus pulposus surrounded by an annulus fibrosis and separated from the vertebral body by a cartilaginous endplate (Humzah and Soames 1988). During one’s lifetime, due to excessive physical loading, occupational injuries, aging, genetics and other factors, the IVDs may degenerate and display marked biochemical and morphological changes (Buckwalter 1995, Urban and Roberts 2003). Currently, magnetic resonance imaging (MRI) is the gold-standard for evaluating disc degeneration. Based on this imaging, numerous methods are available to grade and summarize different features indicative of degeneration, including signal intensity loss, bulging and herniation, as well as disc space narrowing (Battie, Videman, and Parent 2004, Cheung et al. 2009). Lumbar disc degeneration (LDD) is of clinical importance because it is believed to be a major cause of low back pain (Luoma et al. 2000, Livshits et al. 2011, Takatalo et al. 2011, Samartzis et al. 2011). Its severe form lumbar disc herniation (LDH), in which disc material herniates into the epidural space and compresses a lumbar nerve root, can cause neuropathic pains (sciatica) radiating to the lower extremity (Ropper and Zafonte 2015).

Twin studies have demonstrated a strong genetic contribution to LDD (Sambrook, MacGregor, and Spector 1999, Battie et al. 2008). However, searching for genetic variants associated with LDD has been a challenge due to discrepancies and non-standardization of phenotype definitions, inconsistencies with imaging technology, and limited sample sizes in genome-wide association studies (GWASs) (Eskola et al. 2012, Williams et al. 2013, Eskola et al. 2014). Similar to most other complex traits, LDD is likely to be polygenic with thousands of trait-associated variants each of which has tiny effect size.

In addition to age, sex, and environmental influences, LDD is also associated with several heritable risk factors including body mass index (BMI) (Liuke et al. 2005, Samartzis et al. 2011, Samartzis et al. 2012, Takatalo et al. 2013), bone mineral density (BMD) (Harada et al. 1998, Pye et al. 2006, Wang et al. 2011), and serum lipid levels (Leino-Arjas et al. 2008, Longo et al. 2011, Zhang et al. 2016). But it is not fully clear if there is a genetic basis for these phenotype associations. Identifying genetic overlaps between LDD and related traits will be useful for elucidating cause and effect because genetic markers are not subject to reverse causation or confounding and can be used as an instrument to infer causality using Mendelian randomization (Davey Smith and Hemani 2014), and it can also suggest pleiotropic loci that reveal novel insights into biology (Solovieff et al. 2013).

Several methods have been developed to evaluate genetic overlap between traits by exploiting the polygenic architecture (Dudbridge 2016). A polygenic score (PGS) of a trait is the summation of alleles across loci weighted by their effect sizes estimated from GWAS (Purcell et al. 2009). In its typical application, GWAS of a *base phenotype* is first done in a discovery sample. PGS can be calculated in an independent testing sample using SNPs whose p-values are below some threshold in the discovery GWAS. It can then be used as a predictor of *target phenotypes* in the testing sample using regression analysis. PGS has been widely used to predict disease risk (Chatterjee, Shi, and Garcia-Closas 2016), evaluate genetic overlaps across traits (Krapohl et al. 2016), and infer genetic architectures (Stahl et al. 2012, Palla and Dudbridge 2015). Because phenotyping of LDD by MRI is expensive and labor intensive, sample sizes are usually limited for well-phenotyped cohorts. PGS can leverage GWAS meta-analysis results from large consortia to maximize the power to detect genetic overlaps and is most suitable for the current study of LDD. Some other methods, such as bivariate linear mixed-effect model (Lee, Yang, et al. 2012, Vattikuti, Guo, and Chow 2012) would require genotypes of individuals of base and target phenotypes. Recently developed linkage disequilibrium score (LDSC) regression (Bulik-Sullivan et al. 2015) makes use of only summary-level association statistics and can account for sample overlaps between different studies, but it requires very large sample sizes that has not been not available for LDD.

In this study, we applied PGS to investigate the genetic overlap of LDD with four related risk factors using the GWAS data of Hong Kong Disc Degeneration (HKDD) population-based cohort (Cheung et al. 2009, Li et al. 2016, Samartzis et al. 2012) and a Japanese case-control cohort of LDH that required surgery (Song et al. 2013). We selected BMI, BMD and serum lipids levels as base phenotypes, based on their previous reported associations with LDD (Pye et al. 2006, Longo et al. 2011, Samartzis et al. 2012). Height was also included because its association with chronic low back pain (Hershkovich et al. 2013, Heuch et al. 2015). Two semi-quantitative scores that summarized different aspects of LDD from lumbar spine MRI were used as target phenotypes in the HKDD cohort; LDH affection status was used as the third target phenotype in the case-control cohort. Because GWASs of base phenotypes were done in European populations whereas our testing samples were of East Asian ancestry, the performance of PGS in predicting base phenotypes was first evaluated in independent Chinese samples. Then we applied the best performing PGS of the base phenotype to test association with target phenotypes in testing samples (Figure 1). Results were then interpreted in light of previous epidemiological evidence and statistical power to detect association. To better understand the mechanism implied by the genetic overlaps and motivated by the suggestion that LDD and osteoarthritis may share common pathophysiological features (Loughlin 2011, Ikegawa 2013), we further tested if the base phenotypes that had genetic overlaps with LDD also showed genetic correlations with OA using the GWAS summary data of the arcOGEN study (Zeggini et al. 2012). Finally, we evaluated the predictive power of trans-ethnic PGS to aid the design of future studies.

**Figure 1.**
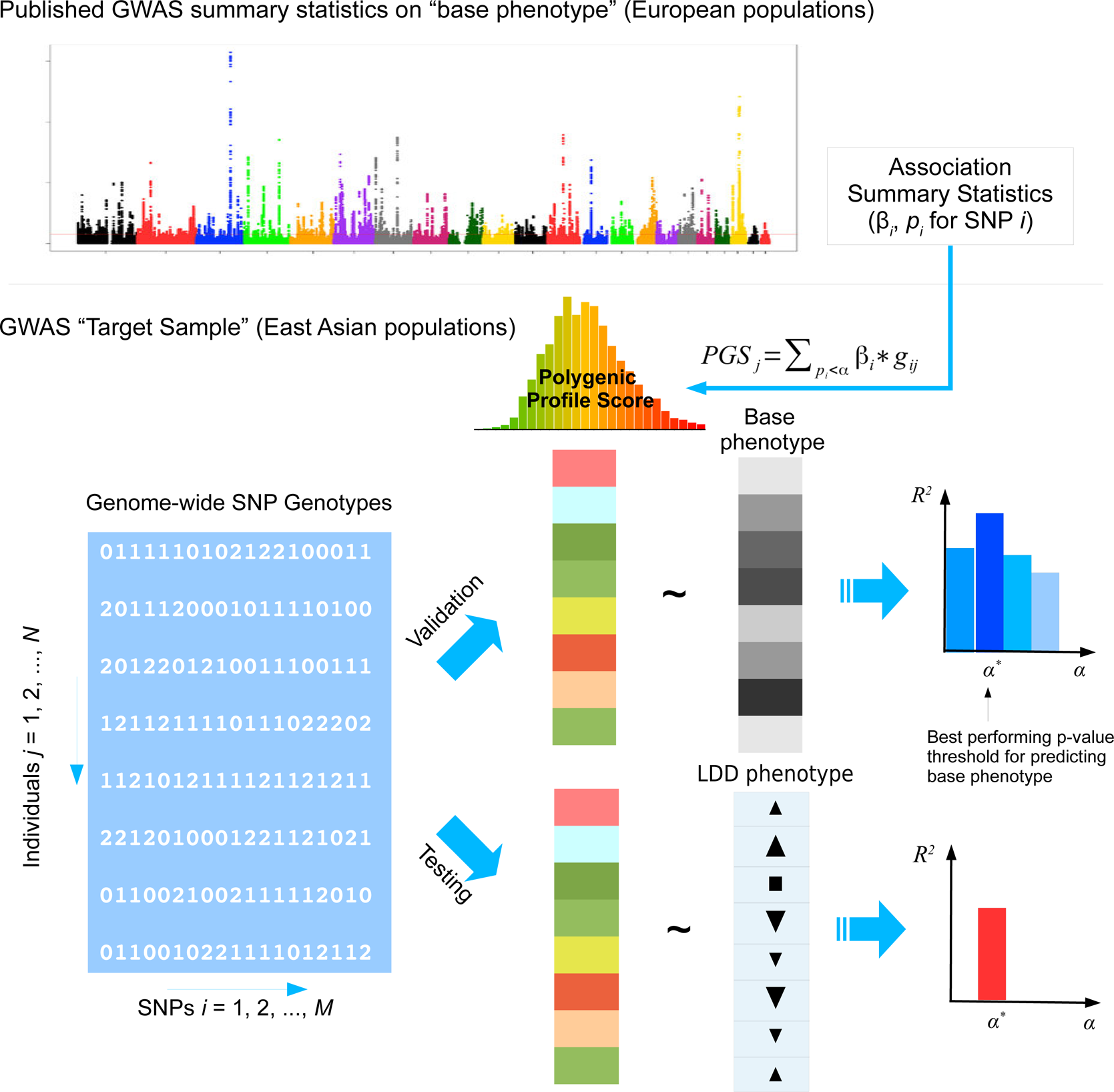
The analysis framework. GWAS summary statistics of base phenotypes were obtained from published studies in European populations. The PGS in a target sample of East Asian population was calculated by weighted summation of alleles at independent SNPs given different p-value thresholds. The performance of PGS in predicting the base phenotype in East Asians was first evaluated in a validation sample. Then the best performing PGS of the base phenotype was used to test association with LDD phenotypes in testing samples.

## Materials and Methods

### Study Samples

#### HKDD

The HKDD Study was a population-based cohort of approximately 3,500 Southern Chinese initiated to assess spinal phenotypes and their risk factors. All participants underwent T2-weighted MRI examination of the lumbar spine assessed by expert physicians (JK and KMC). Sample recruitment and MRI procedures have been described in detail previously (Cheung et al. 2009, Samartzis et al. 2012, Li et al. 2016). For the current study, we focused on two major aspects of LDD captured by different MRI features (Figure 2a). The first was signal intensity loss within nucleus pulposus, which may represent loss of water content of IVD. Its presence and severity at each lumbar disc was assessed by the Schneiderman’s grades (Schneiderman et al. 1987). Based on this grading scheme each disc was given a score of 0 to 3, whereby 0 indicated normal and higher scores indicated increased severity. A *disc degeneration score* for each individual was calculated by the summation of Schneiderman’s grades over all five lumbar discs. We also assessed disc displacement, represented as a bulging/protrusion or extrusion of disc material. An ordinal grade from 0 to 2 was assigned to each lumbar disc to indicate normal, bulge/protrusion or extrusion of disc material; for each individual, a *disc displacement score* was calculated by the summation of the grades over all five lumbar discs (Cheung et al. 2009). Age, sex, physical workload based on occupation, history of smoking, and history of lumbar spine injury were obtained by a questionnaire for all participants. Body height and weight were measured at the time when each subject underwent MRI, and BMI was calculated by dividing weight by height squared (kg/m^2^). A subset of the cohort (*N*=815) also had their blood metabolite profiles measured by quantitative serum nuclear magnetic resonance (NMR) platform (Soininen et al. 2009, Soininen et al. 2015). Low-density lipoprotein cholesterol (LDL-C), high-density lipoprotein cholesterol (HDL-C), triglycerides (TG) and total cholesterol (TC) were obtained as part of NMR metabolite measures. Association between LDD scores and other covariates were analyzed using multiple linear regression to account for correlations between predictor variables. The best fitting model was selected using Akaike information criterion.

**Figure 2.**
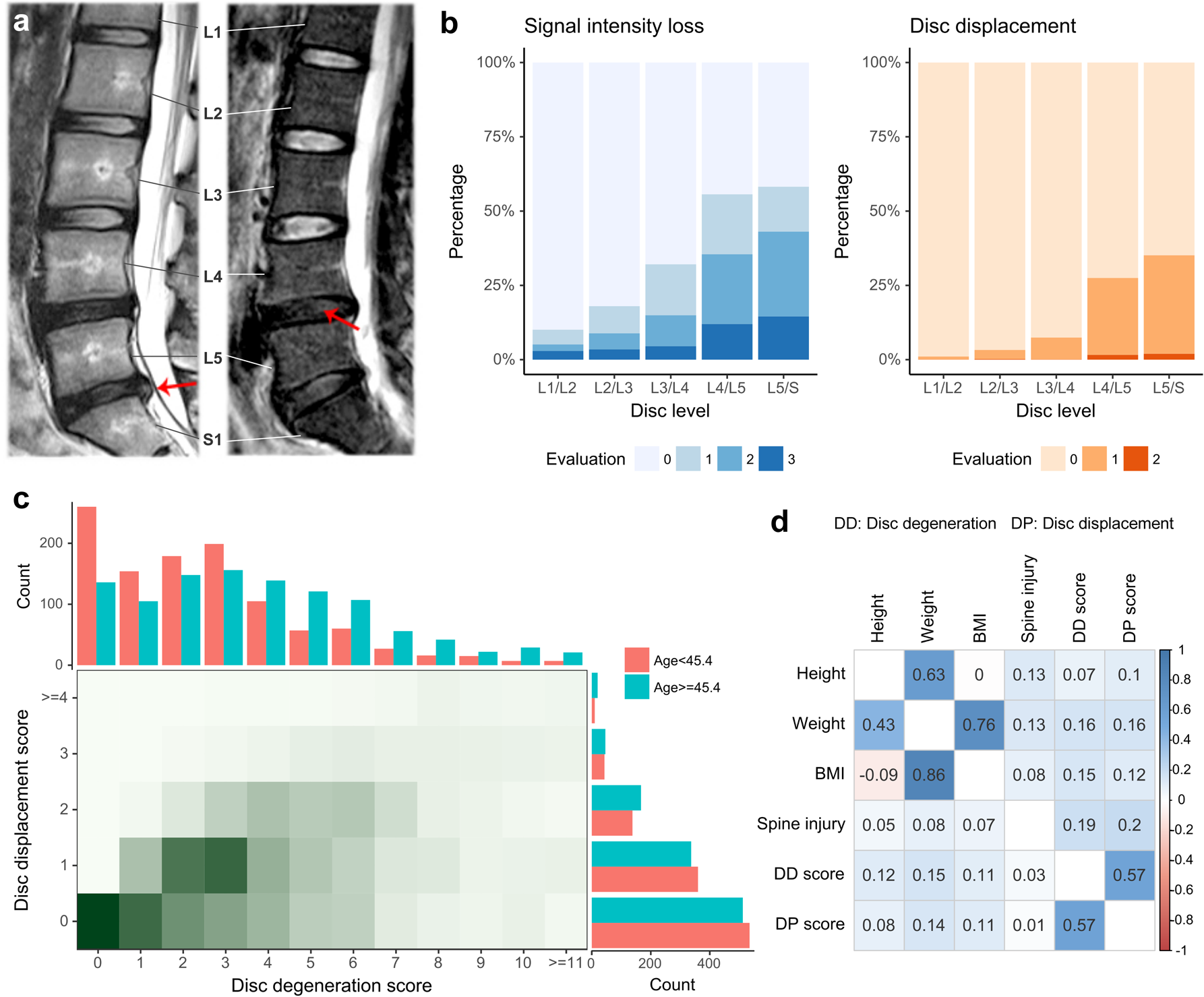
Summary of phenotypes in the HKDD cohort. (a) Examples of MRI images show two major aspects of LDD. Disc displacement (left) is shown as bulging of disc material beyond confine of annulus fibrosus. The loss of proteoglycan and water content (right) within nucleus pulposus is reflected by the signal intensity loss. The lumbar spine has 5 intervertebral segments, termed L1 through L5. S1 stands for the first segment of sacral that is intermediately below the lumbar spine. (b) The prevalence of signal intensity loss and disc displacement at different levels of lumbar spine discs. Two ordinal grades (0-3 for signal intensity loss, 0-2 for disc displacement) were assigned to each lumbar disc to indicate the presence and severity of LDD, where 0 indicated normal and higher scores indicated increased severity. (c) The distribution of LDD scores. The disc degeneration score and displacement score were defined by the summation of grades over all disc levels for signal intensity loss and disc displacement respectively. The two LDD scores are correlated in the population. The age threshold divides the HKDD cohort in two parts with roughly equal sample sizes. The older subjects tend to have higher disc degeneration scores and disc displacement scores. (d) Pairwise Pearson correlations between original phenotypes (upper triangle) and between residual phenotypes after adjusting for age and gender (lower triangle).

A total of 2,373 individuals from the HKDD cohort were genotyped by Illumina HumanOmni-ZhongHua-8 Beadchip. Basic genotyping and quality control procedures have been described in our previous study (Li et al. 2016). In this study, we used more stringent criteria that keep only individuals with a call rate >99% and common SNPs with MAF>0.01. The genotypes were imputed to over 8 million common variants in Phase 3 of 1000 Genomes reference panel using the Michigan Imputation Server (Das et al. 2016) and filtered to keep only common bi-allelic SNPs (MAF>0.01) with imputation quality metrics *r*^2^≥0.3.

#### LDH case-control cohort

The Japanese case-control cohort was part of our previous genetic study of LDD (Song et al. 2013). Hospitalized patients of lumbar disc herniation (LDH) were ascertained on the basis of sciatica or severe low back pain requiring surgical treatment and confirmed by lumbar spine MRI. The controls were unrelated individuals from general Japanese population as part of Japan Biobank Project. All individuals were genotyped by Illumina HumanHap550v3 BeadChip. A total of 366 cases and 3,331 controls passed quality control (QC) and were used in the association analysis. Genotypes were imputed to 2.5 million SNPs in Phase 2 HapMap Project using IMPUTE2 (Howie, Donnelly, and Marchini 2009), and association analysis at each SNP was performed by logistic regression assuming an additive model using SNPTEST (Marchini et al. 2007).

#### HKOS

Hong Kong Osteoporosis Study (HKOS) was a prospective cohort study of over 9,000 Southern Chinese residents in Hong Kong(Cheung, Tan, and Kung 2017). BMD of the lumbar spine (LS-BMD) and femoral neck (FN-BMD) were measured by dual-energy x-ray absorptiometry. The age-corrected and standardized BMD was generated for each gender. A total of 800 unrelated females with extreme BMD were selected in the previous GWAS (Kung et al. 2010). The low BMD subjects were those with BMD Z-score ≤−1.28 at either the LS or FN; high BMD subjects were those with BMD Z-score ≥1.0 at either of the two skeletal sites. All individuals were genotyped by Illumina HumanHap610Quad Beadchip, whereby 780 individuals passed QC. Association analysis at each SNP was performed by linear regression using PLINK (Purcell et al. 2007). Detailed genotyping, QC, and imputation procedures have been described elsewhere (Kung et al. 2010, Xiao et al. 2012).

#### arcOGEN

The arcOGEN study (http://www.arcogen.org.uk/) was a collection of unrelated, UK-based individuals of European ancestry with knee and/or hip osteoarthritis from the arcOGEN Consortium (Panoutsopoulou et al. 2011, Zeggini et al. 2012). Cases were ascertained based on clinical evidence with a need of joint replacement or radiographic evidence of disease (Kellgren–Lawrence grade ≥2), controls were from ancestry-matched (UK) population. A GWAS that included 7,410 cases and 11,009 controls as the discovery sample has been described previously (Zeggini et al. 2012). The summary statistics of the discovery GWAS were obtained by application to the consortium.

All studies were approved by local ethical committees. Written informed consent was obtained from all participants.

### Statistical Analysis

#### Polygenic score regression

We selected height, BMI, BMD, and serum lipid levels as base phenotypes, and obtained GWAS summary data (Table 1). PGS was created by the following two strategies and used to predict phenotypes and test genetic overlap with LDD through linear regression after accounting for other known covariates.

**Table 1.** GWAS summary statistics used in this study.

| Phenotype | Study | Publication | Sample size | Population | Availability |
| --- | --- | --- | --- | --- | --- |
| Height | GIANT Consortium | Wood et al. (2014) | 253,000 | European | Public |
| Body mass index (BMI)<sup>§</sup> |  | Locke et al. (2015) | 234,000 ~ 322,000<sup>#</sup> | European | Public |
| Serum lipids | Global Lipids Genetics Consortium (GLGC) | Willer et al. (2013) | 95,000 ~ 189,000<sup>#</sup> | European | Public |
| Bone mineral density (BMD) | GEFOS2 Consortium | Estrada et al. (2012) | 33,000 | European | Application to the consortium |
|  | Hong Kong Osteoporosis Study (HKOS)<sup>¶</sup> | Kung et al. (2010) | 780 | Chinese | Contributed by collaborators |
| Severe Osteoarthritis | arcOGEN Consortium | Zeggini et al. (2012) | 7,400 cases, 11,000 population controls | European | Application to the consortium |
| Hospitalized lumbar disc herniation | Japan LDH | Song et al. (2013) | 366 cases, 3331 population controls | Japanese | Contributed by collaborators |

<sup>§</sup> The GIANT consortium's BMI GWAS included samples from multiple ethnicities; only the result of the European samples was used.
<sup>#</sup> BMI and lipids summary data were generated from GWAS+Metabochip joint analysis, so sample sizes can vary across different SNPs.
<sup>¶</sup> The HKOS GWAS was part of the GEFOS meta-analysis, and was the only study of non-European population in that study.

The first strategy only used known trait-associated SNPs that reached genome-wide significance in previous studies (GWAS hits). Individual PGS profiles were calculated by summing up the dosage of trait-increasing alleles from imputed genotypes weighted by the reported effect sizes. This strategy has the advantage of including secondary signals within the same locus and increased accuracy of effect size estimates from a larger replication sample. As a second strategy, we performed genome-wide PGS analysis using PRsice (Euesden, Lewis, and O’Reilly 2015). Briefly, summary statistics of base GWAS was first aligned with genotyped SNPs of the testing sample. Then SNPs were pruned based on p-value informed clumping algorithm (LD *r*^2^<0.1 across 250kb) that selected SNPs most associated with the base phenotype in a locus to generate sets of independent SNPs. PGS was created using clumped SNPs whose p-value in the base GWAS were below pre-specified threshold. We varied p-value thresholds (1.0E-7, 1.0E-5, and from 1.0E-4 to 0.5 with a step of 0.0001) to select the one that maximized the variance of explained (*R*^2^) for the base phenotype in the validation sample. When individual-level genotype data in the validation sample were not available (the HKOS GWAS and LDH case-control cohort), PGS regression was performed using summary statistics based algorithm implemented in gtx R package (Johnson 2012), and SNP genotypes of HapMap 3 East Asian samples were used as the reference panel for SNP clumping.

#### Correcting sample overlap and extreme selection

The HKOS GWAS was part of BMD GWAS meta-analysis conducted by the GEFOS consortium. To make it an independent testing data, we inverted the fixed effect meta-analysis to subtract the contribution of HKOS from the GEFOS summary statistics (Supplementary Note 1). When calculating PGS using GWAS hits, we used effect estimates from the stage II replication sample to avoid the issue of overlapping sample.

The HKOS GWAS adopted an extreme phenotype design to increase association power, which also resulted in upward biased estimates of *R*^2^ by PGS using linear regression. To get an estimate of *R*^2^ in the unselected sample (*R̂*^2^) for comparing with the previous report, we corrected the *R*^2^ by the PGS of known BMD-associated SNPs in the selected sample (*R̂*^2^′):

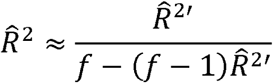

where *f* is the increased phenotype variance due to extreme phenotype selection (=2.739 in the HKOS GWAS sample). The derivation and validation of this approximation formula is given in Supplementary Note 3.

#### *R*^2^ for case-control data on the liability scale

For the case-control data, it is meaningful to estimate the disease liability explained by PGS under the liability threshold model (Falconer and Mackay 1996), so that the result can be compared to the heritability of LDH (Heikkila et al. 1989). We first converted summary statistics generated by the logistic regression to those of linear regression by first-order approximation. Then we used summary statistics based PGS regression to obtain an estimate of *R*^2^ on the observed scale. Finally, the observed *R*^2^ was converted to the liability scale using the transformation formula by Lee, Goddard, et al. (2012) assuming the disease prevalence of 0.02 (Jordan, Konstantinou, and O’Dowd 2009). More details are given in Supplementary Note 4.

#### Inference of genetic architecture and projecting prediction performance

We applied AVENGEME (Palla and Dudbridge 2015) to estimate parameters of genetic architecture of height and BMI from the PGS results. The procedure is described in Supplementary Note 2. For a presumed SNP heritability 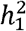, the method estimated the fraction of markers that are null (*π̂*_0_) and genetic correlation between the discovery and testing samples (*σ̂*_12_). If the genetic architectures of the discovery and testing sample are the same, then the genetic correlation between two sample can be estimated as 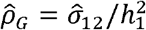.

The same model was also applied to predict the expected *R*^2^ for height and BMI in Chinese population under different study designs. To project *R*^2^ using PGS created by weights from European GWAS, model parameters were set to the maximum likelihood estimates fitted to the observed PGS results. We also increased the discovery sample size by 500,000 to evaluate the increase of *R*^2^ in the future. To predict *R*^2^ using PGS created by weights from East Asian GWAS, we used the same set of model parameters, but set *N*_1_ to the sample sizes of largest GWAS of East Asians (He et al. 2015, Wen et al. 2014) and assumed no heterogeneity of effect sizes between the discovery and testing samples (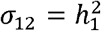). To incorporate between-sample heterogeneity within East Asians, we changed between population genetic correlation to 0.9 (so *σ*_12_ = 0.9*h*^2^), which was a lower bound for height and BMI in Europeans (de Vlaming et al. 2017) and in different Chinese GWAS samples (data not shown).

#### SNP heritabilities

Phenotypes analyzed in the HKDD cohort were adjusted by known covariates; and residues were inverse normal transformed when necessary. SNP heritability of the adjusted phenotypes were estimated using GCTA v1.25 (Yang, Lee, et al. 2011) after excluding individuals so that no pair of individuals had estimated coefficient of relatedness >0.0275.

#### Estimating genetic correlation between anthropometric traits and osteoarthritis

To test the genetic overlaps osteoarthritis with height, BMI and BMD, we applied LDSC regression (Bulik-Sullivan et al. 2015) to the GWAS summary statistics following the recommended procedure. BMD summary statistics were corrected to remove the contribution from the HKOS GWAS (the only non-European study) as in PGS regression.

Due to sample overlaps in different European GWASs, PGS could not have been applied to the genome-wide summary data. But for known BMD associated SNPs whose effect size estimates from an independent replication sample were available, we also used PGS regression under summary statistic mode to assess the genetic correlation between osteoarthritis and BMD.

#### Power analysis

Power calculation was done assuming the test statistics follows non-central chi-squared distribution under the alternative hypothesis. The non-centrality parameter for quantitative trait is 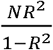, where *N* is the sample size and *R*^2^ is the phenotype variance explained by PGS. For binary trait, *R*^2^ in the above formula is on the observed scale and can be converted from liability scale using Lee, Goddard, et al. (2012)’s formula as described in Supplementary Note 4.

## Results

### Phenotype summary of the HKDD cohort

A total of 2054 unrelated Chinese subjects in the HKDD cohort (60% were females) were included in polygenic analysis. The basic demographic and phenotype summary are shown in Table S1. Both signal intensity loss and disc displacement showed a higher prevalence and severity at lower lumbar levels (Figure 2b). The disc degeneration and disc displacement scores for each individual were calculated by the summation of grades over all levels. Consistent with a major effect of aging, older individuals tend to have higher disc degeneration and displacement scores (Figure 2c). The two LDD scores were highly correlated (*r* =0.57; Figure 2c). Both of them were also positively correlated with height, body weight, BMI, and lumbar spine injury (*P*<0.001; Figure 2d, Table S2a); the correlation remained significant for all except injury after correcting for the effect of age and gender (Figure 2d, Table S2b). Multiple linear regression analysis showed that the best fitting models for both disc degeneration and displacement scores included age, sex, lumbar injury, height and BMI as covariates (Table S3), which together explained 21.5% and 9.6% phenotype variances respectively. The SNP heritability estimates for height and BMI in the HKDD cohort were 0.38 (±0.18) and 0.25 (±0.17), similar to the previous reports in Europeans (Yang et al. 2010, Yang, Manolio, et al. 2011). For disc degeneration and displacement scores, after adjusting for known covariates, SNP heritability estimates were about 0.2~0.3 (Table 2).

**Table 2.** SNP heritability estimates of phenotypes analyzed in the HKDD cohort.

| Phenotype | Adjustment and transformation | $\hat{h}^2$ (SE) | *p*-value |
| --- | --- | --- | --- |
| Height | Age, Sex;<br>inverse normal transformation | 0.533 (0.170) | 6.75E-05 |
|  | Age, Sex, First two PCs;<br>inverse normal transform | 0.383 (0.182) | 1.67E-02 |
| BMI | Age, Age<sup>2</sup>, Sex;<br>inverse normal transformation | 0.285 (0.171) | 2.97E-02 |
|  | Age, Age<sup>2</sup>, Sex, First PC;<br>inverse normal transformation | 0.249 (0.174) | 6.03E-02 |
| Disc<br>degeneration<br>score | Age, Sex, Lumbar Injury | 0.218 (0.163) | 6.50E-02 |
|  | Age, Sex, Lumbar Injury, Height | 0.232 (0.169) | 6.43E-02 |
|  | Age, Sex, Lumbar Injury, BMI | 0.226 (0.170) | 7.26E-02 |
|  | Age, Sex, Lumbar Injury, Height,<br>BMI | 0.219 (0.171) | 8.40E-02 |
|  | Age, Sex, Lumbar Injury, Weight | 0.225 (0.171) | 7.80E-02 |
| Disc<br>displacement<br>score | Age, Sex, Lumbar Injury | 0.291 (0.176) | 4.26E-02 |
|  | Age, Sex, Lumbar Injury, Height | 0.269 (0.180) | 6.20E-02 |
|  | Age, Sex, Lumbar Injury, BMI | 0.238 (0.181) | 9.17E-02 |
|  | Age, Sex, Lumbar Injury, Height,<br>BMI | 0.216 (0.182) | 1.21E-01 |
|  | Age, Sex, Lumbar Injury, Weight | 0.213 (0.182) | 1.22E-01 |

### Evaluating the prediction performance of polygenic scores of anthropometric traits

We first evaluated prediction performance of PGS in Chinese samples and compare them with Europeans. PGS profiles of height and BMI were created in the HKDD cohort using known trait-associated SNPs identified by the GIANT consortium GWAS meta-analyses. They explained 5.7% and 1.2% of height and BMI variances respectively, after adjusting for age, sex and principle components. It is 2~3 fold lower than previous reports in independent European samples, which were 16% for height (Wood et al. 2014) and 2.7% for BMI (Locke et al. 2015). The prediction performance of BMD associated SNPs reported by GEFOS consortium were tested in the HKOS GWAS sample (Kung et al. 2010). After correcting for extreme phenotype selection (Supplementary Note 3), the known BMD-associated SNPs explained 3.4% and 3.0% variance of LS-BMD and FN-BMD in Chinese population, also lower than previous reported ~5% in Europeans (Estrada et al. 2012).

Since GWAS hits may only explain a small proportion of phenotype variance, we extended PGS analysis to make use of whole-genome summary statistics (Figure 3). As the p-value threshold of the discovery GWAS increases, both true and false positive SNPs will be included in the PGS. The p-value threshold that optimized phenotype prediction depends on the discovery sample size and unknown genetic architecture (Chatterjee et al. 2013, Dudbridge 2013), and should be determined empirically. At the optimal p-value threshold, we found the phenotype variance explained is similar to that using GWAS hits for FN-BMD, marginally improved for height, slightly worse for LS-BMD, and more than doubled for BMI (2.6%). Theoretical model fitting under a range of plausible parameters (Table S4) suggested that BMD had smaller fraction of trait associated SNPs (0.5~0.6%) with larger effect sizes than BMI and height (estimated fraction of non-null markers: 14~17%), which explained why sparse PGS models showed better prediction performance for BMD. The trait variances explained by PGS predicted by the models generally captures the trend of empirical observations at different p-value thresholds (Figure 3). The estimate of between-population genetic covariance for each phenotype was consistently lower than the presumed heritability (Table S4), reflecting trans-ethnic heterogeneity in effect sizes.

**Figure 3.**
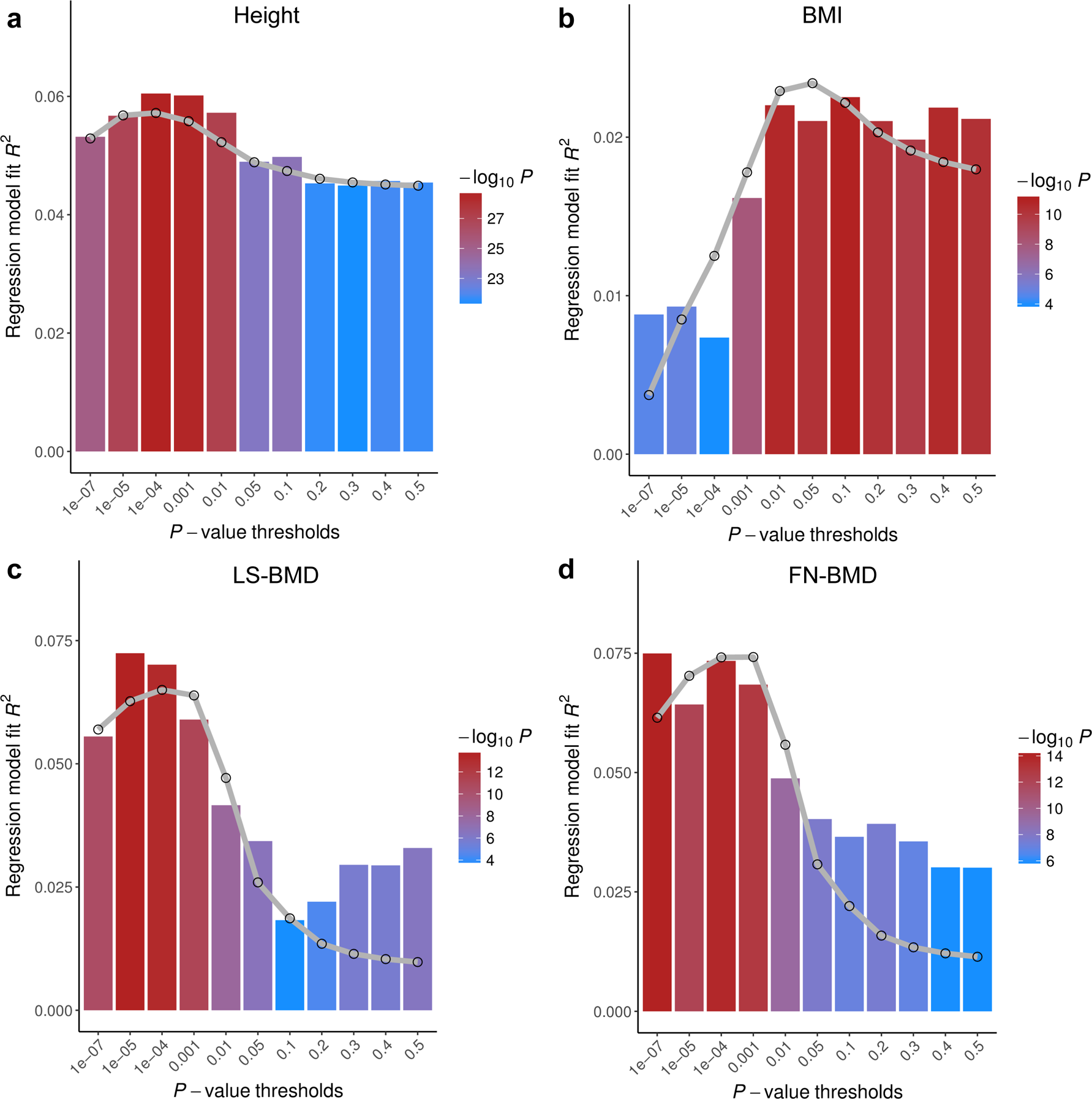
Prediction performance of polygenic scores (PGS) on four base phenotypes in Chinese population. Phenotype variances explained (*R*^2^) are shown at different p-value thresholds. The gray lines are the predicted *R*^2^ based on the theoretical model of Dudbridge (2013) with parameters given in Table S4. Different parameter set of each model give similar results. PGS of height and BMI are tested on HKDD cohort; PGS of bone mineral density at the lumbar spine (LS-BMD) and femoral neck (FN-BMD) are evaluated on HKOS sample.

### Testing genetic overlaps between anthropometric traits and LDD

We then applied PGS to test genetic overlaps between anthropometric traits and LDD (Table 3). In the HKDD cohort, the BMI PGS at its optimal threshold was positively associated with both disc displacement score (*R*^2^=0.24%, *P*=0.025) and disc degeneration score (*R*^2^=0.23%, *P*=0.028) after adjusting for sex, age and lumbar injury. The results are consistent with obesity as a major risk factor for LDD development and progression (Hangai et al. 2008, Hassett et al. 2003). The associations remained significant (*P*<0.05) after further adjusting for height but disappeared after adjusting for BMI or body weight (Table S5). The PGS of LS-BMD were positively associated with disc displacement score (*R*^2^≈0.2%; *P*<0.05) and remained significant (*P*≤0.05) after further adjusting for height, BMI or weight (Table S6). The same trend was also observed for FN-BMD but did not reach significance. The finding supports the previous reported genetic correlation between BMD and disc bulge in a twin study (Livshits et al. 2010). The lack of association with disc degeneration score is also consistent with the previous study that showed a smaller effect size between BMD and disc signal intensity on MRI (Livshits et al. 2010).

**Table 3.** Genetic overlap of lumbar disc degeneration with anthropometric traits. For the four base phenotypes, we adopted two strategies to create polygenic profiles to predict target phenotypes: using known trait-associated SNPs and their effect size estimates, or using independent SNPs of GWAS summary statistics selected based on the optimal p-value threshold for predicting the base phenotype.

| Base Phenotype | SNP Predictor | $N(\text{GWAS Sample})^b$ | $N(\text{SNPs})^c$ | $R^{2d}$ | Association with disc displacement score<sup>e</sup> | | | Association with disc degeneration score<sup>**</sup> | | | Association with lumbar disc herniation requiring surgery | | |
| --- | --- | --- | --- | --- | --- | --- | --- | --- | --- | --- | --- | --- | --- |
| | | | | | $\text{sgn}(\beta)$ | $R^2$ | $p\text{-value}$ | $\text{sgn}(\beta)$ | $R^2$ | $p\text{-value}$ | $\text{sgn}(\beta)$ | $R^{2f}$ | $p\text{-value}$ |
| Height (Wood et al. 2014) | Known loci | 253,000 | 622 | 5.76% | + | 0.003% | 8.04E-01 | + | 0.135% | 9.40E-02 | + | 0.351% | 7.73E-03 |
| | PGS $P \leq 2.0\text{E-}04$ | | 4093 | 6.63% | + | 0.022% | 5.02E-01 | + | 0.147% | 7.98E-02 | + | **0.339%** | **8.92E-03** |
| BMI (Locke et al. 2015) | Known loci | 234,000 ~ 322,000 | 92 | 1.24% | + | 0.016% | 5.46E-01 | + | 0.053% | 2.94E-01 | - | 0.000% | 9.80E-01 |
| | PGS $P \leq 1.23\text{E-}02$ | | 4359 | 2.64% | + | **0.240%** | **2.53E-02** | + | **0.230%** | **2.84E-02** | + | 0.011% | 6.37E-01 |
| Lumbar Spine BMD (Estrada et al. 2012) | Known loci | 46,000 | 60 | 9.33%<sup>g</sup> | + | **0.222%** | **3.16E-02** | + | 0.000% | 9.92E-01 | + | 0.093% | 1.71E-01 |
| | PGS $P \leq 1.0\text{E-}05$ | 32,000 | 110 | 7.25% | + | 0.197% | 4.27E-02 | + | 0.007% | 7.07E-01 | + | 0.128% | 1.08E-01 |
| Femoral Neck BMD (Estrada et al. 2012) | Known loci | 51,000 | 60 | 7.78%<sup>g</sup> | + | 0.062% | 2.55E-01 | - | 0.017% | 5.51E-01 | + | 0.073% | 2.24E-01 |
| | PGS $P \leq 7.0\text{E-}04$ | 33,000 | 578 | 7.97% | + | 0.147% | 8.01E-02 | + | 0.036% | 3.86E-01 | + | 0.182% | 5.51E-02 |

<sup>b</sup> Sample size of discovery sample GWAS. For known BMD-associated loci, shown are the sample size of second stage replication cohort.
<sup>c</sup> Number of SNPs in the HKDD cohort that passed QC and have minor allele frequency $\geq 0.01$ .
<sup>d</sup> Variance of base phenotype in Chinese population explained by the PGS. For height and BMI, $R^2$ was estimated in the HKDD cohort ( $N=2054$ ). Height was adjusted for age, sex, and the first two principle components (PCs); BMI was adjusted for sex, age, age<sup>2</sup>, and the first PC. The residuals were then inverse normal transformed. For BMD, it was estimated in the HKOS GWAS sample (Kung et al. 2010; $N=780$ females). LS-BMD and FN-BMD were adjusted for age and standardized into Z-scores.
<sup>e</sup> Association with disc displacement and degeneration scores were evaluated by inclusion of polygenic profile score as a covariate to the multiple linear regression model of target phenotype that adjust for age, sex and lumbar spine injury.
<sup>f</sup> For LDH, $R^2$ is the variance of disease liability explained by the PGS (Supplementary Note 4).
<sup>g</sup> HKOS sample were selected from extreme ends of BMD distribution. After correcting for extreme-selection (Supplementary Note 3), we estimate that $R^2$ by the PGS of GWAS hits is 3.52% for LS-BMD and 2.99% for FN-BMD.

**Table 4.**
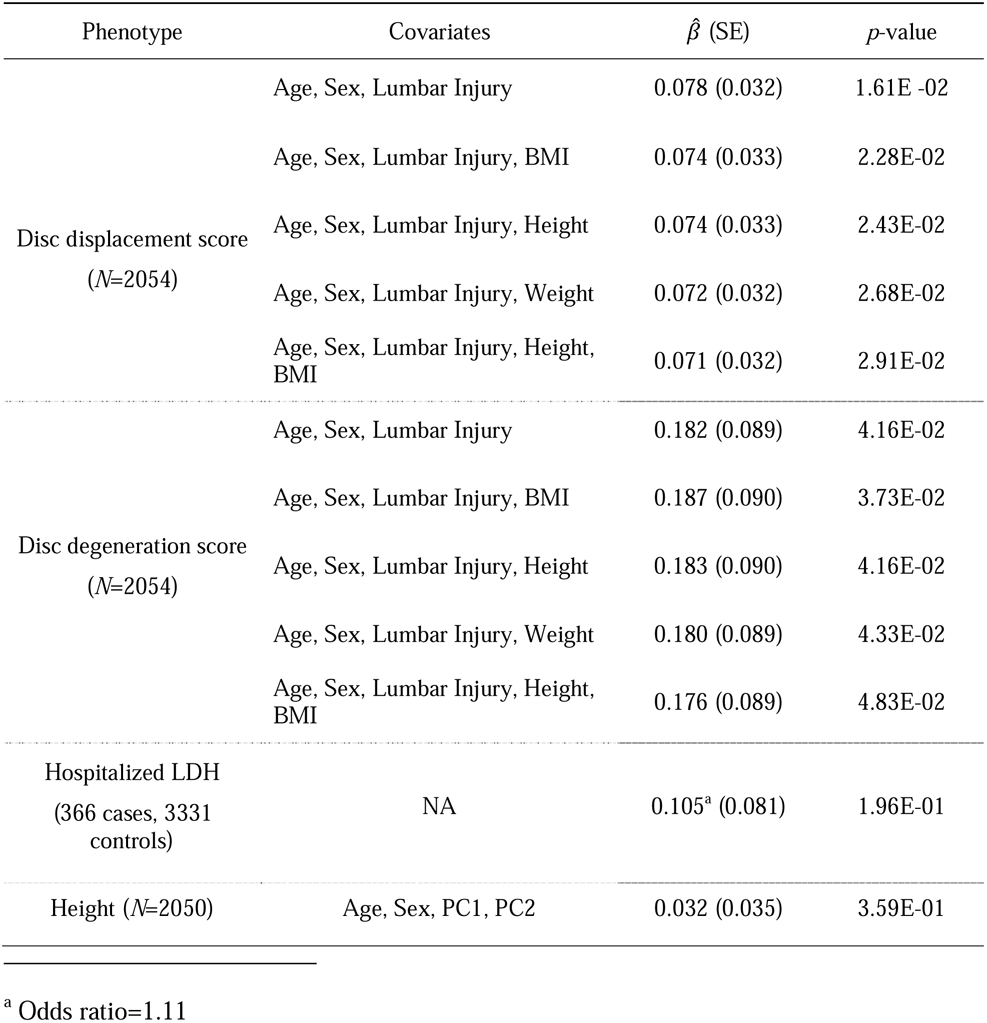
Association of rs4733724-A allele with lumbar disc degeneration and height in East Asian samples. The SNP rs4733724 was genotyped in the HKDD cohort and reliably imputed in the Japanese case-control cohort. The A allele was previously reported to be associated with increased height in Europeans (Wood et al. 2014). The rs4733724-A allele was coupled to rs6651255-T, the latter of which was recently found to increase the risk (odds ratio=1.23) of LDH requiring surgery in Icelanders (Bjornsdottir et al. 2017). The frequency of rs4733724-A allele is 0.72 in East Asians and 0.23 in Europeans.

| Phenotype | Covariates | $\hat{\beta}$ (SE) | *p*-value |
| --- | --- | --- | --- |
| Disc displacement score<br>(*N*=2054) | Age, Sex, Lumbar Injury | 0.078 (0.032) | 1.61E-02 |
|  | Age, Sex, Lumbar Injury, BMI | 0.074 (0.033) | 2.28E-02 |
|  | Age, Sex, Lumbar Injury, Height | 0.074 (0.033) | 2.43E-02 |
|  | Age, Sex, Lumbar Injury, Weight | 0.072 (0.032) | 2.68E-02 |
|  | Age, Sex, Lumbar Injury, Height, BMI | 0.071 (0.032) | 2.91E-02 |
| Disc degeneration score<br>(*N*=2054) | Age, Sex, Lumbar Injury | 0.182 (0.089) | 4.16E-02 |
|  | Age, Sex, Lumbar Injury, BMI | 0.187 (0.090) | 3.73E-02 |
|  | Age, Sex, Lumbar Injury, Height | 0.183 (0.090) | 4.16E-02 |
|  | Age, Sex, Lumbar Injury, Weight | 0.180 (0.089) | 4.33E-02 |
|  | Age, Sex, Lumbar Injury, Height, BMI | 0.176 (0.089) | 4.83E-02 |
| Hospitalized LDH<br>(366 cases, 3331 controls) | NA | 0.105<sup>a</sup> (0.081) | 1.96E-01 |
| Height (*N*=2050) | Age, Sex, PC1, PC2 | 0.032 (0.035) | 3.59E-01 |

<sup>a</sup> Odds ratio=1.11

**Table 5.** Genetic correlations estimated by LD-score regression.

| Trait 1 | Trait 2 | $\hat{r}_G$ (SE) | $p$-value |
| --- | --- | --- | --- |
| *Genetic correlations between anthropometric traits* |  |  |  |
| Height | BMI | -0.055 (0.022) | 1.36E-02 |
| Height | Lumbar spine BMD | 0.071 (0.032) | 2.72E-02 |
| Height | Femoral neck BMD | 0.036 (0.033) | 2.83E-01 |
| BMI | Lumbar spine BMD | 0.067 (0.028) | 1.75E-02 |
| BMI | Femoral neck BMD | 0.071 (0.028) | 9.90E-03 |
| Lumbar spine BMD | Femoral neck BMD | 0.669 (0.032) | 4.64E-96 |
| *Genetic correlations between osteoarthritis and anthropometric traits* |  |  |  |
| Osteoarthritis | BMI | 0.255 (0.050) | 4.02E-07 |
| Osteoarthritis | Height | 0.117 (0.045) | 9.50E-03 |
| Osteoarthritis | Lumbar spine BMD | 0.192 (0.076) | 1.18E-02 |
| Osteoarthritis | Femoral neck BMD | 0.094 (0.068) | 1.63E-01 |

In addition to LDD scores in the general population, we also applied PGS to predict case-control status of symptomatic LDH (Song et al. 2013). The height PGS was positively associated with LDH (*P*<0.01) and explained 0.35% of disease liability. The association cannot be explained by body weight, because the BMI PGS is better associated with weight but does not show association with LDH (*P*>0.5). The result provides a genetic basis to the previous epidemiological observation that being tall is a risk factor for hospitalization due to LDH (Wahlstrom et al. 2012) and back surgery (Coeuret-Pellicer et al. 2010).

### Testing genetic overlaps between lipid levels and LDD

Previous studies also reported that increased level of LDL-C, TC and TG were associated with increased risk of LDH (Longo et al. 2011, Zhang et al. 2016, Leino-Arjas et al. 2008). To test if serum lipid levels have genetic correlation with LDD, we did similar PGS analysis using known lipid associated SNPs and GWAS summary data from the Global Lipids Genetic Consortium (Willer et al. 2013). Prediction performance of PGS was first evaluated in a subset of the HKDD cohort (*N*=620 with genotypes) whose lipid levels were measured by the high-throughput NMR approach. All PGS were significantly associated with the corresponding lipid levels. Except for LDL-C, the PGS of known lipid loci showed the best prediction performance (Figure S1). However, none of them was significantly associated with LDD scores with the expected direction in the HKDD cohort or LDH in the Japanese case-control cohort (Table S7). Directly testing the phenotype association in the HKDD cohort by multiple linear regression also showed no association (Table S8). Therefore, our data does not support the previously suggested role of atherosclerotic lipids in LDD.

### Power consideration

Given sample sizes and study designs, the two testing samples used in this study show similar profiles of statistical power (Figure 4). We have >50% power (at significance level *α*=0.05) to detect genetic correlation if PGS explains >0.2% variance of adjusted LDD scores (or LDH disease liability). To achieve the same power at *α*=0.01, it would require PGS to explain >0.33% phenotype (liability) variance. Therefore, the current study does not have enough power to detect genetic overlap if the PGS explain less than 0.2% variance of LDD scores (liability). To reduce the multiple testing burden, we had only used the PGS which was optimal in predicting the corresponding base phenotype to test the association with LDD. The results that were nominally significant and consistent with the expected phenotype correlations can be interpreted as *supportive evidence to genetic overlaps*.

**Figure 4.**
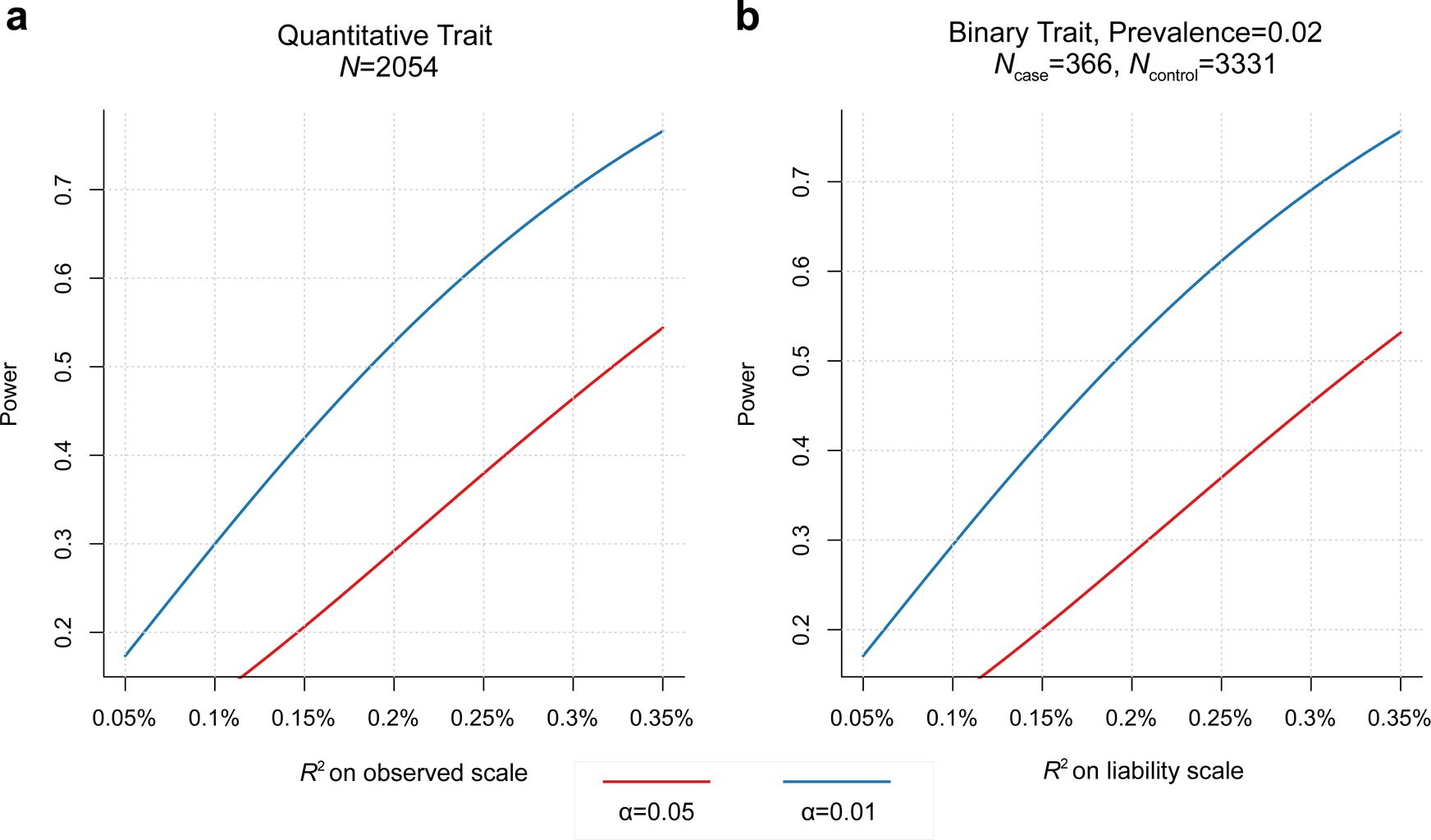
Power to detect association with polygenic score (PGS) in two testing samples. (a) For the HKDD cohort (*N*=2054), given significance level (at *α*=0.01 or 0.05), the power is determined by the phenotype variance explained by PGS. (b) For the Japanese case-control cohort (366 cases, 3331 controls), assuming the disease prevalence of 0.02, the power is a function of the disease liability explained by PGS.

**Figure 5.**
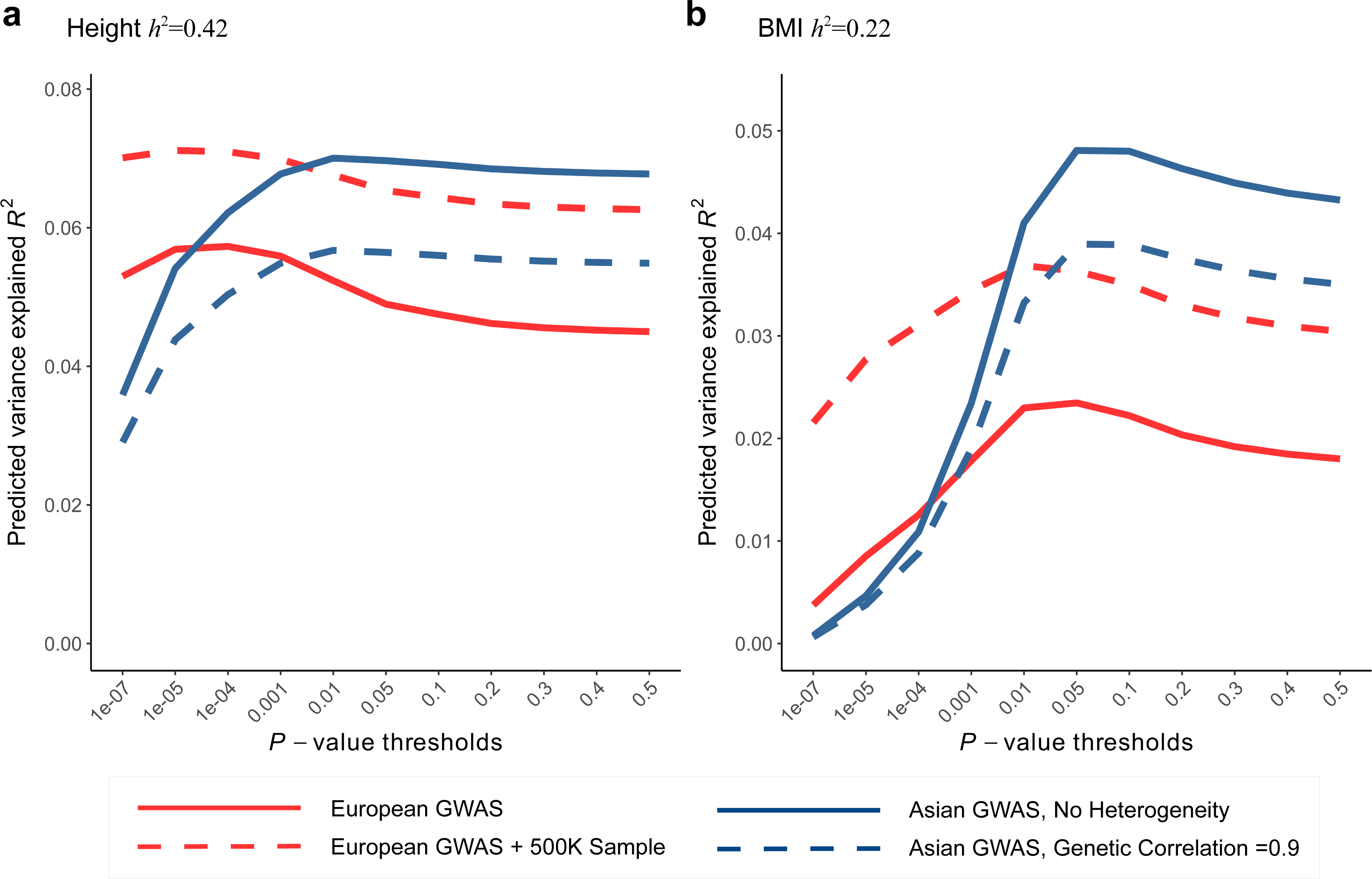
Comparing the prediction performance of polygenic score (PGS) in Chinese sample using discovery GWAS of European and East Asian. Expected phenotype variances explained by PGS (*R*^2^) were calculated using the theoretical model of Dudbridge (2013) with parameters compatible with the observed PGS results of height and BMI. (a) For height, the latest European GWAS has sample size 252K. Assuming SNP heritability of 0.42 in both populations, expected *R*^2^ in Chinese population (red line) was calculated using the parameters best fit to Figure 3. In comparison, the latest East Asian GWAS has sample size only 36K, expected *R*^2^ was calculated using the same set of parameters except that we assumed no heterogeneity in effect sizes (i.e., genetic correlation=1) between discovery and testing sample (blue line). To predict the gain in *R*^2^ when using even larger European GWAS in the future, we further increased the discovery GWAS sample size by 500K (red dashed line). We also relaxed the assumption of no heterogeneity within East Asian and calculate expected *R*^2^ assuming genetic correlation of 0.9 (blue dashed line). (b) For BMI, European GWAS has sample size 234K, East Asian GWAS has sample size 87K. Expected *R*^2^ were calculated similarly as height, assuming SNP heritability of 0.22.

### The association of a height associated SNP rs6651255 with LDD scores

The current study had no power to search for individual SNPs showing pleiotropic associations with LDD and other traits. But we noted that a recent GWAS of LDH with lumbar spine surgery in Iceland population (Bjornsdottir et al. 2017) identified a genome-wide significant SNP rs6651255, which was also a known height associated SNP (Wood et al. 2014). The risk allele T showed an odds ratio of 1.23 and associated with increased height. The same study also reported that increase in the genetically determined height increased the risk of LDH with surgery, but the effect of rs6651255 on LDH was not mediated by height. To replicate this finding in our cohorts, we found an LD proxy rs4733724 (LD *r*^2^=1 with rs6651255 in 1000 Genomes CEU population) was directly genotyped in the HKDD cohort and reliably imputed in the Japanese case-control cohort. The allele A was coupled to the LDH risk allele and significantly increased both disc displacement and disc degeneration scores (*P*<0.05; Table 3). The effects remained significant after further adjusting for height, BMI or body weight. The same allele was also weakly associated with increased height and increased risk of LDH requiring surgery, but the results were not significant as the sample sizes limited the power to detect small effect sizes.

### Genetic correlation between anthropometric trait and osteoarthritis

Finally, LDD has been suggested to share common features with osteoarthritis which is also known as degenerative joint disease (Ikegawa 2013; Loughlin 2011). To test if osteoarthritis also showed genetic overlaps with the same set of traits as LDD, we assessed the genetic correlations of BMI, BMD and height with osteoarthritis using LDSC regression (Table 2). Significant positive genetic correlation was found between BMI and osteoarthritis (*r̂*_G_ = 0.255, *P*=4.0E-07), which is expected given the strong evidence for a causal role of BMI (Panoutsopoulou et al. 2014). Suggestive positive genetic correlations with osteoarthritis were also observed for height (*P*=9.5E-03) and LS-BMD (*P*=0.012) but not for FN-BMD (*P*>0.1).

The genetic correlation between osteoarthritis and LS-BMD was less significant though its effect was stronger than between osteoarthritis and height, which was possibly due to smaller sample size of the BMD GWAS. Since the genetic architecture of BMD was dominated by fewer number of causal SNPs with larger effect sizes (Table S4), it is also possible that LDSC which assumed an infinitesimal model may be less optimal to detect genetic correlations for BMD and other traits. To support this, we calculated PGS of BMD GWAS hits using weights from the second stage replication sample of the GEFOS consortium (Estrada et al. 2012) to predict osteoarthritis. The PGS of both LS-BMD and FN-BMD were strongly associated with osteoarthritis case-control status in the acrOGEN sample (*R*^2^=0.13%, *P*=7.8E-07 for LS-BMD and *R*^2^ =0.12%, *P*=2.5E-06 for FN-BMD). Taken together, the results suggest that like LDD, osteoarthritis also shares genetic overlaps with height, BMI and BMD.

## Discussion

### Between-population heterogeneity and its impact on prediction performance of polygenic score

In this study, we adopted a trans-ethnic PGS strategy to evaluate the genetic overlaps between different traits where GWAS of base phenotypes were done in Europeans and validation and testing samples were East Asians. Although most GWAS findings were generally replicated in populations different from the initial discovery, heterogeneity commonly existed in the estimated effect sizes (e.g., Marigorta and Navarro 2013, Carlson et al. 2013), which would reduce the power of PGS to predict phenotypes in populations from a different ethnicity (e.g., Johnson et al. 2015). We also found in Chinese validation samples that variance of height, BMI and BMD explained by the PGS of corresponding GWAS hits were consistently lower than in Europeans. For height and BMI assuming the genetic architecture is the same in European and Chinese, the observed PGS results are consistent with between-population genetic correlations about 0.4~0.6 (Table S4, Materials and Methods). The rough estimations are within the range of previous estimate for type 2 diabetes and rheumatoid arthritis between European and East Asian using a different methodology (Brown et al. 2016).

The use of European GWAS in the current study is mainly due to large sample sizes and publicly available summary statistics. GWAS meta-analyses of height and BMI were also conducted in East Asians (He et al. 2015, Wen et al. 2014). Although sample sizes are much smaller (*N*≈36,000 for height, 87,000 for BMI), they are expected to have more similar effect sizes to the Chinese sample. To evaluate the tradeoff between sample size and effects heterogeneity, we projected expected prediction performance (*R*^2^) of height and BMI using a theoretical model with parameters of genetic architecture compatible with the observed PGS results (Materials and Methods). Despite smaller sample sizes, using East Asian GWAS as the discovery sample is expected have comparable maximum *R*^2^ as European GWAS to predict height in the Chinese population (Figure 3). For BMI, depending on the presumed SNP heritability, using East Asian GWAS shows comparable or better maximum *R*^2^ (Figure 3, Figure S2). Further increase the European GWAS sample sizes by half a million, a scale similar to the on-going UK biobank study, the increase in *R*^2^ for height is capped at 8% but roughly doubles for BMI. Notably, when using East Asian GWAS as the discovery sample, the best prediction performance can only be achieved at p-value thresholds >0.01. However, whole-genome summary statistics of East Asian GWASs are not publicly available for us to take advantage of the power gain. Also consistent with the theoretical predictions, incorporating East Asian GWAS top hits or adding new height association SNPs from a recent exome-chip study (Marouli et al. 2017) to the PGS of GWAS hits only marginally increased *R*^2^ in predicting height and BMI, and their associations with LDD scores remained insignificant (Table S9).

Given genetic architecture and sample sizes, the power of PGS in detecting genetic overlaps is mainly determined by the performance PGS in predicting the corresponding base phenotype. Therefore, the theoretical results suggest that the use of European GWAS as discovery sample in PGS analysis can still be a favorable approach in cross-trait analysis in the East Asian population. But we noted that when applying PGS in a testing sample of different ethnicity from the discovery GWAS make it not straight forward to estimate or interpret genetic correlations between different traits. We also caution that the trans-ethnic PGS strategy may not be suitable for other populations like African. Nevertheless, whenever possible ancestry-matched GWAS of base phenotype with large sample sizes should be used to improve the power and interpretability of the results.

### The influence of phenotype definition

In this study, we analyzed three LDD phenotypes, including two semi-quantitative scores derived from MRI assessment and one clinically defined symptom. The PGS of height, BMI and BMD were associated with at least one LDD phenotype. It highlights the complexity in operationally defining LDD, as the current diagnostic approach only captures certain aspects of the degenerative process. Therefore, comparison between different studies should clarify how phenotypes are defined. And it will be fruitful to jointly evaluate multiple MRI features in future genetic studies. However, although MRI is the current gold standard that gives best resolution in defining LDD, it is too expensive to be carried out in large samples.

An alternative strategy is to use the a “proxy phenotype” such as patient-based LDH in which large number of cases can be identified based on electronic medical records. Use of proxy phenotype has been demonstrated to improve the power in GWAS (e.g., Okbay et al. 2016). Increase in sample sizes has been shown outweigh the dilution of genetic effects, but it may also capture certain aspects of the trait that is irrelevant to the phenotype of interest (e.g., Kong et al. 2017). In the current and our previous study (Song et al. 2013), LDH requiring surgery was presumed to represent an extreme end of disc displacement in the population. In this regard, it is surprising that the PGS of height strongly associated with LDH but not LDD scores, and PGS of BMI and BMD were associated with LDD scores but not LDH. Although the lack of expected associations can be false negatives due to insufficient power, we cannot rule out the possibility that ascertainment of LDH patients based on severe low back pain or sciatica may enrich polygenic factors other than LDD.

### Biological interpretations

The observed genetic overlaps can be explained by either causality or genetic pleiotropy or both. Interestingly, BMI, BMD and height also showed suggestive evidence of positive genetic correlation with osteoarthritis. It is possible that they can be explained by some common mechanisms. Although formal assessment of causality could utilize the Mendelian randomization paradigm although this requires larger sample sizes, PGS can be used to nominate candidate phenotypes (Evans et al. 2013). Overweight or obesity has been established as one of the major risk factors for the development and progression of both LDD (Hassett et al. 2003, Hangai et al. 2008) and osteoarthritis (Bierma-Zeinstra and Koes 2007). It is commonly believed that increased body weight or BMI exerts more physical loading to the IVD and vertebral endplate (Videman, Levalahti, and Battie 2007) or joint cartilage (Guilak 2011), and leads to increased wear and tear of the structures. For BMD, in addition to its correlation with LDD, previous studies also found the increase in BMD in osteoarthritis patients and an inverse association between osteoarthritis and osteoporosis (Hannan et al. 1993, Arden et al. 1996). It was postulated that increased BMD is associated with a loss of resilience of subchondral bone which may results in increased mechanical stress on joint cartilage (Foss and Byers 1972, Radin and Rose 1986) and similarly on IVD (Harada et al. 1998). The causal mechanism of tall stature on LDH that leads to hospitalization or surgery remains unclear. One possibility may be related to increased disc height, because a previous study using finite element modeling demonstrated that discs with taller height and smaller area were prone to larger motion, higher annular fiber stress and larger degree of disc displacement (Natarajan and Andersson 1999). Another possibility may be altered spinal alignment in taller individuals that predispose them to lumbar spine injury. Notably, the postulated mechanisms all point to the pathophysiological role of biomechanical stress. Some other mechanisms have also been proposed (Katz, Agrawal, and Velasquez 2010, Samartzis et al. 2013). For example, obesity is also believed to lead to local inflammatory response of secondary mediators secreted by adipocytes known as adipokines. The causal role of adipokines and inflammatory markers can also be tested using their genetic predictors as instrument in future studies.

Alternatively, the observed genetic correlations are also consistent with the genetic pleiotropy and shared pathways among skeletal phenotypes. In supporting this notion, several individual osteoporosis associated SNPs were associated with height or BMD (Reynard and Loughlin 2013), and osteoporosis and LDD were found to share some common genetic risk factors (Song et al. 2008, Williams et al. 2011). At single SNP level, we also replicated the recent finding of Bjornsdottir et al. (2017) and showed that the height-associated SNP rs6651255 was associated with two LDD scores. The previous study did not find association of the same SNP with other related skeletal phenotypes like osteoarthritis of the spine or osteoporotic vertebral fractures and suggested that the association was driven by the neuropathic pain rather than herniated lumbar discs. However, they did not examine the association of the SNP with radiologically defined LDD phenotypes. Our results in the large population-based cohort with MRI assessment suggest that the same SNP also influences the changes in composition and morphology of lumbar discs. Future genetic studies on LDD with larger sample sizes should search for additional pleiotropic SNPs to better understand bone-cartilage relationships.

In summary, the current study is the first attempt to evaluate genetic overlap between LDD and related traits using GWAS data. Our trans-ethnic polygenic analysis supports the genetic correlations of height, BMI and BMD with LDD, and sheds new light on understanding the pathological mechanism of degenerative skeletal disorders.

## Acknowledgements

This work was supported by Research Grant Council of Hong Kong Theme-based Research Scheme “Functional Analyses of How Genomic Variation Affect Personal Risk for Degenerative Skeletal Disorders” (T12-708/12N). GEFOS study was funded by the European Commission (HEALTH-F2-2008-201865-GEFOS). arcOGEN study was funded by a special purpose grant from Arthritis Research UK (grant 18030). We thank Ms. Pei Yu for curating the HKDD phenotype database, and Dr. Eleftheria Zeggini for providing the arcOGEN GWAS summary data.

## Author Contributions

XZ and PCS conceived the study and coordinated the research. XZ developed the methods and performed analysis. JK and KMC evaluated MRI images of the HKDD cohort. TK, YS, KC, YK and SI contributed the Japanese LDH GWAS summary data. CLC contributed the HKOS GWAS summary data. YH contributed the GEFOS consortium GWAS summary data. DS contributed the NMR data. YL and DC applied for the arcOGEN consortium GWAS summary data. XZ drafted the manuscript with inputs from PCS, KSC. DS, TMS, CLC and JK reviewed and revised manuscript. PCS and KCS supervised the study.

